# Evaluating the data quality of iNaturalist termite records

**DOI:** 10.1101/863688

**Authors:** Hartwig H. Hochmair, Rudolf H. Scheffrahn, Mathieu Basille, Matthew Boone

**Author notes:** Corresponding author (HH).

## Abstract

Citizen science (CS) contributes to the combined knowledge about species distributions, which is a critical foundation in the studies of invasive species, biological conservation, and response to climatic change. In this study, we assessed the value of CS for termites worldwide. First, we compared the abundance and species diversity of geo-tagged termite records in iNaturalist to that of the University of Florida termite collection (UFTC) and the Global Biodiversity Information Facility (GBIF). Second, we quantified how the combination of these data sources affected the number of genera that satisfy data requirements for ecological niche modeling. Third, we assessed the taxonomic correctness of iNaturalist termite records in the Americas at the genus and family level through expert review based on photo identification. Results showed that iNaturalist records were less abundant than those in UFTC and in GBIF, although they complemented the latter two in selected world regions. A combination of GBIF and UFTC led to a significant increase in the number of termite genera satisfying the abundance criterion for niche modeling compared to either of those two sources alone, whereas adding iNaturalist observations as a third source only had a moderate effect on the number of termite genera satisfying that criterion. Although research grade observations in iNaturalist require a community-supported and agreed upon ID below the family taxonomic rank, our results indicated that iNaturalist data do not exhibit a higher taxonomic classification accuracy when they are designated research grade. This means that non-research grade observations can be used to more completely map the presence of termite locations in certain geographic locations without significantly jeopardizing data quality. We concluded that CS termite observation records can, to some extent, complement expert termite collections in terms of geographic coverage and species diversity. Based on recent data contribution patterns in CS data, the role of CS termite contributions is expected to grow significantly in the near future.

## Introduction

Termites are destructive insect pests that annually cause billions of dollars in damage and losses. These costs, based on insecticide sales figures, were estimated to be $22 billion worldwide in 1999 [1]. Using 2010 sales data, the estimated global economic impact of termites has increased to $40 billion, with subterranean termites accounting for about 80 % of the costs [2]. Understanding spatio-temporal patterns of termite distributions is therefore necessary for the implementation of effective pest control strategies. Open-access databases provide unprecedented access to biodiversity data in the form of species occurrences worldwide. Biodiversity databases vary in composition of data sources, and include expert surveys, georeferenced museum collection data, published literature reviews, and contributions from recreational naturalists. Specifically, the continued advancement of information and communication technology, including the emergence of the Web 2.0 and the development of location-aware mobile technologies, sensors, and cameras have greatly increased the capacity of how citizens can contribute to Citizen Science (CS) projects [3]. Citizens have become an important source of geographic information, even in domains that had until recently been the exclusive realm of authoritative agencies [4]. However, because of their cryptic nature and small size, termite specimens retrieved from their substrata or during dispersal flights often require expert knowledge [5], which renders data collection and identification particularly challenging. Previous research has demonstrated that CS adds scientific value to conventional science in various aspects, including greater geographic scales and temporal range, improved field detection, and detection of unusual occurrences [3]. It also offers an additional way to monitor Essential Biodiversity Variables (EBVs) [6] that cannot be remotely sensed, such as taxonomic diversity or migratory behavior. Furthermore, CS adds other benefits to conservation efforts, natural resource management, and environmental protection through public engagement [3]. The development of spatial statistical methods has led to the frequent use of open-access biodiversity databases in species distribution modeling (SDM) [7], which may be used for a variety of purposes, such as identifying species diversity hotspots [8] and predicting potential ranges of invasive species [9], in addition to forecasting effects of climate change on biodiversity [10]. Consistent survey coverage along time, space, and environment is required to answer different ecological and evolutionary questions and to develop accurate SDMs [11].

iNaturalist is a prominent CS platform for documenting species observations across the world. Participants submit media (pictures, video, or audio) of biological sightings to the iNaturalist data portal that are then identified online by the iNaturalist community [12]. Despite their strong contributions to publicly available data sets, CS based geo-data collections oftentimes suffer from geographic and user selection bias caused by the opportunistic nature of the data collection process [13] compared to data from administrative agencies [14].

While there is no reference data collection available that captures the presence of termites in all parts of the world, it is nevertheless possible to compare various features (e.g. number of sightings, spatial coverage, species diversity) between different online data portals and expert collections. The first task of this study was thus to compare global record numbers, spatial coverage, and species diversity for termites between iNaturalist, the University of Florida termite collection (UFTC) and Global Biodiversity Information Facility (GBIF). The latter data source is currently the largest occurrence data portal, combining data from many sources of different countries, scientific institutions and CS platforms. Since iNaturalist is one the sources feeding into the GBIF platform, its contribution for termite data records in GBIF can be quantified. The second task was to assess how many genera recorded in individual or combined data sources, respectively, satisfy data requirements for ecological niche modeling. The third task was to analyze the taxonomic correctness at the genus and family level of termite records on iNaturalist through photo identification, and to examine if iNaturalist research grade records are more likely to have fewer taxonomic misclassifications than records of other data quality assessment levels.

## Materials and methods

### UF termite collection

The UFTC at the Fort Lauderdale Research and Education Center in Davie, Florida, was established in 1985. An online repository [15] hosts over 38,000 termite samples with official species names from collections that are preserved in 85 % ethanol. Samples were primarily collected by operatives in the pest control industry, property owners, and academics and submitted to the second author (RHS) for identification. The collection database records describe genus and species, geographic latitude and longitude, collection date, and type of structure infested. It contains records from all continents, although the geographic focus is on North and South America. Worldwide, the collection dates range between 1915 and 2019. As of mid 2019 the UFTC contains samples from over 800 unique bionomials.

### iNaturalist

iNaturalist was created in 2008 by graduate students at the University of California Berkley, and is currently funded by the California Academy of Sciences (starting 2014) and the National Geographic Society (starting 2017). Observations from iNaturalist frequently include date, time, and a media record of the observation. Any media records are then identified by the online community. As of 2019, iNaturalist contained over 24 million biological observations from users around the world. Termite records were downloaded from iNaturalist on 4/16/2019 using the R [16] package “rinat” [17] and the query ‘taxon_name = “Termites” ‘. This returns all observations of the epifamily Termidoidae (formerly known as Infraorder Isoptera). After removal of records with missing or invalid coordinates a total set of 6078 termite observations remained worldwide. iNaturalist identifies the reliability of observations through a process called data quality assessment (DQA) [18]. The DQA addresses to some extent all elements of standardized principles for describing the data quality for geographic data [19], which include completeness, logical consistency, positional accuracy, temporal accuracy, and thematic accuracy. The DQA categorizes contributed data into three categories, which are “Needs ID”, “Research Grade” and “Casual”. If an observation satisfies a set of specific technical criteria (i.e. having a date, geographic coordinates, photos or sounds, and not being a captive or cultivated organism), this observation is considered verifiable, and is labeled “Needs ID”, otherwise it is called “Casual”. An observation reaches research grade status (the highest level) if the community agrees at a level lower than family, which is when more than two thirds of two identifiers or more agree on a taxon. From the extracted sample of 6078 termite observations, the vast majority are still “Needs ID” observations (5161 records, 84.9 %), and only 785 (12.9 %) are research grade, with the highest proportion (33.9 %) in South Africa. That latter fact can be partially attributed to the more than 300 termite contributions in that region by user Tony Rebelo, a conservation biologist from the South African National Biodiversity Institute. The remaining records of the worldwide collection on iNaturalist are of “Casual” quality with 132 records (2.2 %). These numbers illustrate that using research grade only records for any type of spatial analysis seriously affects data abundance in difficult to ID taxa.

By default, all photos uploaded to iNaturalist are released under a Creative Commons (CC) Attribution-Non-Commercial (CC-BY-NC) license, which allows free use of the data for non-commercial purposes, as long as the owner is cited. However, users can revoke this license entirely or chose from different versions of the CC license, such as CC0 (essentially public domain, under which data are made available for any use without restriction), or CC BY (under which data are made available for any use, including commercial purposes, provided that attribution is given appropriately for the sources of data used, in the manner specified by the owner).

### GBIF

The GBIF platform is an inter-governmental, global research infrastructure that facilitates the sharing, discovery and access to primary biodiversity data [20]. It provides free access to digitized biological data from different sources (e.g. museum collections, survey programs) as a result of collaborations between data providers and taxonomists across many institutions. As of fall 2019, GBIF facilitated access to over 1.35 billion records per the GBIF website. It gathers data records from more than 800 institutions worldwide, where CS programs represent about 27% of total GBIF records for insects [12]. Data publishers can use a variety of tools, protocols and standards to publish primary occurrence records to GBIF.

Access to records on GBIF can be achieved through bulk download from the GBIF website, through web services, or through third-party libraries, such as the R “rgbif” package [21]. For this study, a GBIF bulk download for geo-tagged records including human observation, material sample or specimen of the Blattodea order was conducted on 4/18/19, followed by application of the filter for the nine termite families of the epifamiliy Termitoidae. This process resulted in 37,061 records worldwide. This dataset contained 443 contributions from three CS programs (iNaturalist: *n* = 436, naturgucker: *n* = 4, natusfera: *n* = 3). Besides this, 35,520 records stem from non-CS sources and 1,098 records lack a data source description. Overall, the share of CS based contributions of termite data to GBIF is small (∼1.2 %). A large bulk of GBIF termite data comes from a single contributor, namely the Commonwealth Scientific and Industrial Research Organisation (CSIRO) with 14,134 records, almost all of which were acquired in Australia.

All occurrence records indexed on GBIF need to satisfy certain data quality criteria before being included in the platform [22]. One of these criteria is that records need to be associated with one of the three CC licenses mentioned before (CC0, CC BY, CC BY-NC), and be of research grade quality. For example, from among the set of 785 worldwide iNaturalist research grade termite records, only 437 observations have one of these three CC licenses. This explains that the number iNaturalist termite records extracted through GBIF (*n* = 436) matches (closely) that for records downloaded directly from iNaturalist when the CC and research grade filters are applied (*n* = 437). The difference of one record stems from the fact that GBIF misses the most recent iNaturalist termite record (dated 4/13/2019) that may not have been discovered in the latest crawling process.

### Mapping termite record distribution

To analyze the distribution of termite records we imported the cleaned data sets into R 3.6.1. We created 2.5° hexagon grid cells for mapping distribution using the st_make_grid function and summarized the number of termite observations in each data set using the sf_intersect function in the SF package [23].

### Data suitability for niche modeling

Ecological niche modeling techniques, such as Maxent, require a sufficient volume of geo-referenced occurrence records with temporal attributes [20]. The recommended minimum number of distinct data-points for niche modeling analysis falls between 10 and 20 [24]. We determined the number termite genera that feature enough observations for ecological niche modeling in any of the analyzed data sources and their combinations, respectively. For this purpose, the surface of the Earth is subdivided into 0.1 degree grid cells [20]. Next, the number of those termite genera is determined which are present in at least 10, 20, and 50 distinct 0.1 grid cells around the globe, respectively. Only records with a time stamp containing at least month and year are considered for this purpose.

### Taxon correctness of iNaturalist termite records

To determine taxon correctness, we performed a manual review of 2201 iNaturalist records in the Americas that have been named at the family rank or lower. The correctness of order, family, subfamily and genus entries for these records is evaluated based on manual inspection of community provided photographs conducted by RHS. The analysis shows how frequently designations at these different taxonomic ranks were misclassified. The choice to conduct the evaluation at the genus rank or higher is because species are often hard to recognize from photographs. Such effort would require samples to be viewed under a microscope. Furthermore, on a subset of 1792 iNaturalist records that were identified at the genus or species taxonomic rank, we determined if taxon correctness of genus and family level depended on research grade status of the observations using a randomization test for contingency tables. This limitation to genus and species taxonomic rank was chosen since, in order to reach research grade status, an observation in iNaturalist must be below family rank. The geographic focus of the termite taxonomic accuracy assessment was restricted to the Americas because of taxonomic expertise of RHS for termites in that part of the world.

## Results

### Data abundance

Contributions to GBIF, iNaturalist, and the UFTC between 1920 and 2018 showed contrasted results (Fig 1). Annual contributions to the UFTC grew steadily starting from the mid 1980’s when RHS established the database, began annual field excursions in central and South America for data collection, and promoted the collection in the pest control industry. After peaking in 2013, the number of annual contributions was surpassed by iNaturalist in 2016. Unlike other submission sources, GBIF expert contributions were sporadic throughout our time period with punctuated uploads between 1995-1999 and 2010-2014. The spike in observations from 1995-1999 was primarily associated with 6489 preserved specimens locations from the Yucatan peninsula in Mexico in 1997. These contributions were administered by the Comisión Nacional para el Conocimiento y Uso de la Biodiversidad (MBM-UACAM). They were collected over the entire year and taken at multiple locations in natural areas. This suggests that the source of this data was actual field data from concerted collection efforts rather than bulk upload from archived data. Another notable peak in expert GBIF contributions fell within the 2010-2014 time range. It was caused by 7249 contributions of the Laboratory of Applied Entomology of the University of Lome which were collected in rural areas of northern Ghana and Togo within approximately three weeks in August 2013. For the GBIF annual charts, it must be noted that over 4500 records (specimen collections) lacked a contribution year, and that these were therefore missing for the GBIF charts but present in contribution maps. The GBIF charts make clear that CS based contributions to GBIF only started in recent years and make up only a small fraction of all GBIF data as of now. The increase in contributions from iNaturalist follows the website’s inception in 2008 and shows that the bulk of data contributions is of non-research grade. This highlights the importance of assessing the taxonomic classification accuracy for those kind of records.

**Fig 1.**
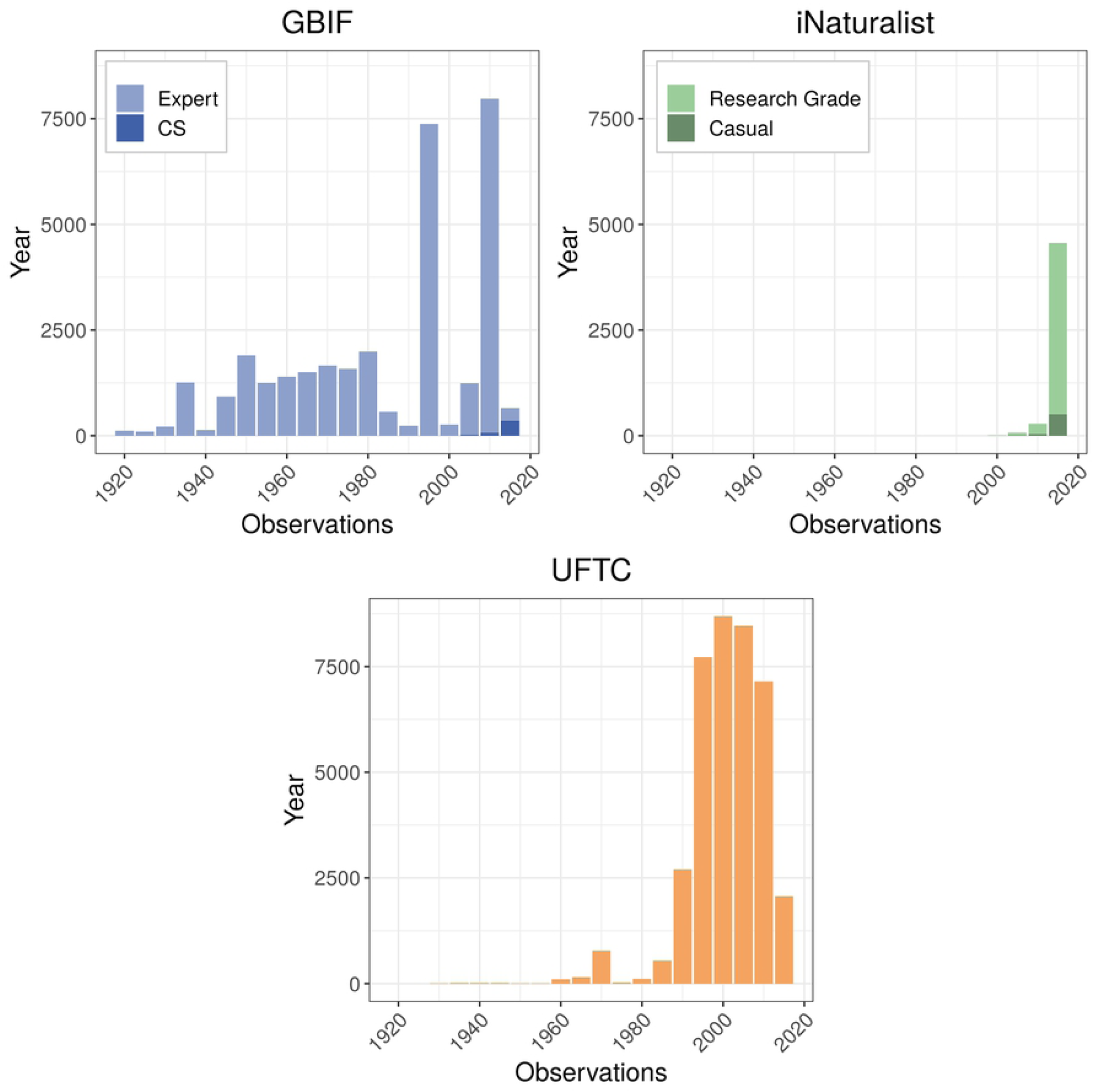
5-year counts of termite records contributed to GBIF, iNaturalist, and the UFTC across the globe between 1920 and 2018.

Australia (including Oceania) was the continent with most expert (non-CS) contributions to GBIF, which stem primarily from CSIRO records mentioned above (Table 1). The iNaturalist community contributed termite observations most actively in North America. Since GBIF CS data stem primarily from iNaturalist, this also resulted in GBIF CS data abundance being highest in North America. Most data records in the UFTC can be found in North America, followed by South America due to frequent field expeditions by RHS.

**Table 1.** Number of termite records in GBIF, iNaturalist, and the UFTC, split by continent. Highlighted cells show the continent with the highest collection count per data category.

| Continent | GBIF (*n* = 37061) |  |  | iNaturalist (*n* = 6078) |  | UFTC (*n* = 38558) |
| --- | --- | --- | --- | --- | --- | --- |
|  | Expert | CS | Unknown | Research | Non-research |  |
| Africa | 7329 | 111 | 27 | 263 | 677 | 2607 |
| Asia | 320 | 53 | 457 | 81 | 417 | 267 |
| Australia | 16047 | 15 | 283 | 18 | 337 | 294 |
| Europe | 489 | 15 | 138 | 24 | 73 | 24 |
| North America | 10250 | 247 | 48 | 394 | 3441 | 27272 |
| South America | 1085 | 2 | 145 | 5 | 348 | 8094 |
| Total | 35520 | 443 | 1098 | 785 | 5293 | 38558 |

Visual inspection of worldwide distribution maps suggests that GBIF termite records were most abundant in Australia (Fig 2a), whereas iNaturalist and UFTC records were mostly found in North America (Fig2b and c).

**Fig 2:**
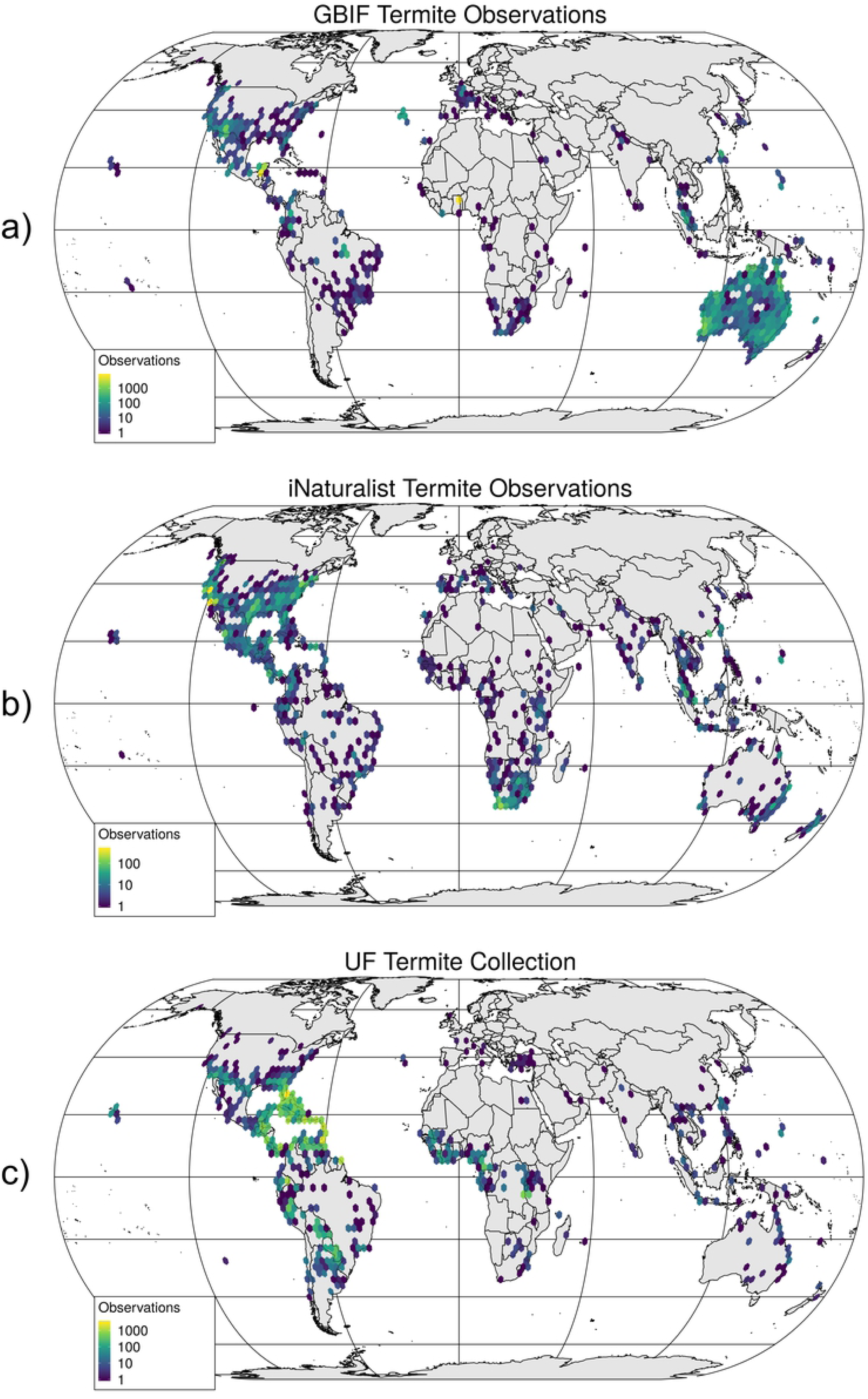
Geographic distribution of termite records in GBIF (a), iNaturalist (b), and the UFTC (c).

The world map in Fig 3a reveals that CS (all sources, including both research and non-research grade iNaturalist records) collected data cover significant observation gaps of expert data in most continents, including regions, such as Southern and Eastern Africa, Southern Europe, India, South-East Asia and New Zealand. This means that CS data are important for the improvement of worldwide termite distribution models.

**Fig 3:**
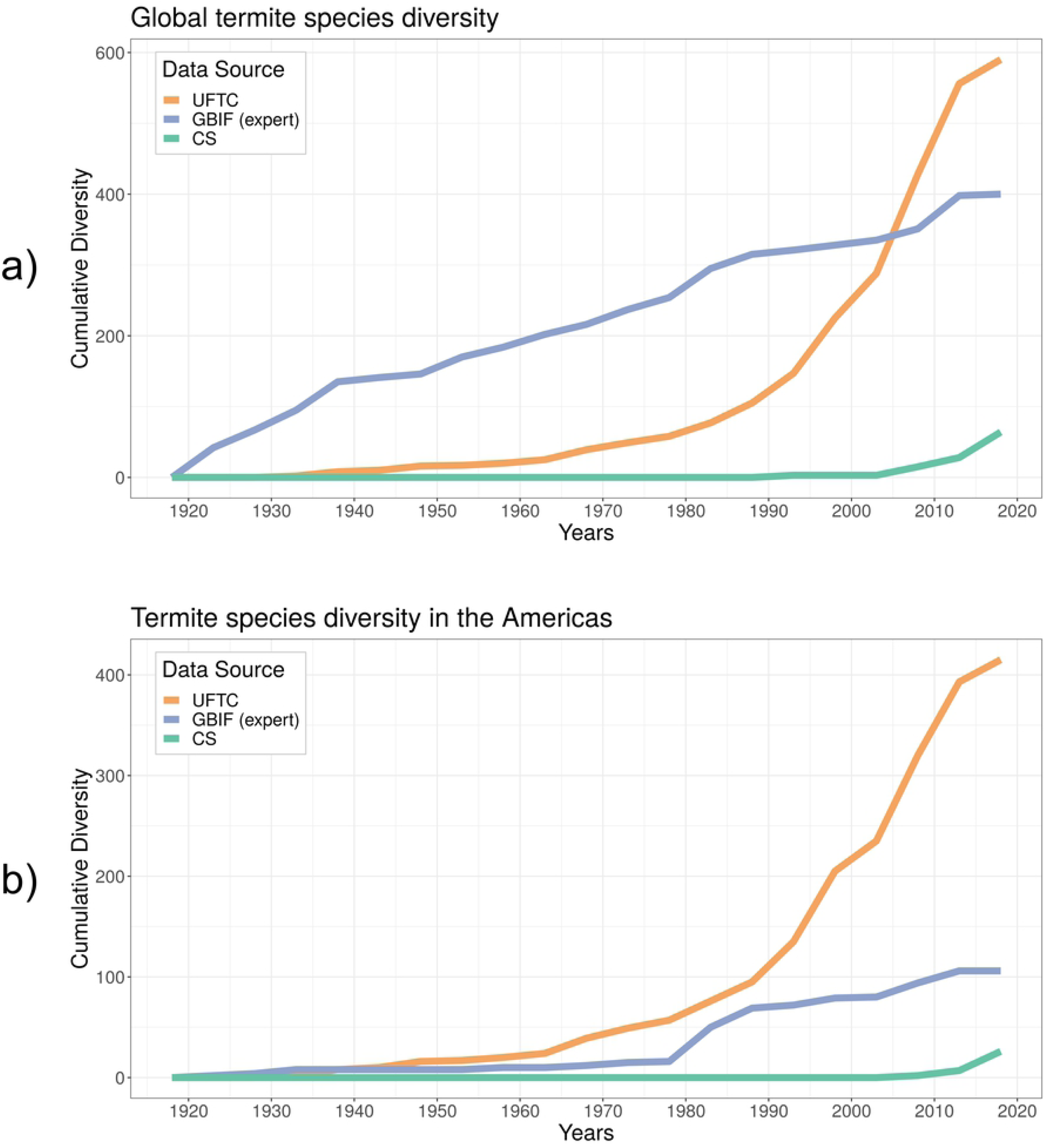
Coverage of termite observations from the three analyzed sources separated into CS based and expert based (a), and coverage of iNaturalist records alone, separated by observation quality (b).

The geographic distribution of termite collection localities for iNaturalist in Fig 3b clearly demonstrates that presence of these data in most world regions through this CS source relies on non-research grade observations (Casual or Needs ID). Comparison to Fig a also shows that several world regions that are not covered by expert users are covered by casual observations in iNaturalist, e.g. in Southern Europe, coastal Brazil, India, or South-East Asia. Given that no difference in termite classification accuracy between iNaturalist research grade and non-research grade observations could be found, it is suggested that considering non-research grade CS observations, especially in combination with expert databases, helps to paint a more comprehensive and clearer picture of global biodiversity than expert or research-grade observations only.

### Species diversity

Cumulative species diversity over the years contrasted expert observations (UFTC, GBIF without CS sources), from CS sources (i.e., GBIF CS contributions and iNaturalist records regardless of quality grade) (Fig 4a). In any of the sources only a small fraction of termite species was collected each year compared to the total of 655 species estimated to be established in the Americas alone [25]. This means that annual termite collection activities miss a large portion of termite species and their spatial coverage. However, since termites spread slowly, i.e. typically less than 100 m/year except for anthropogenic dispersal [26], this problem can be somewhat mitigated by aggregating infestation records over a multi-year time period. Although the total number of records was comparable between GBIF and UFTC globally (compare Table 1), global species diversity was more pronounced in UFTC. Since UFTC contains more than three times the number of records for North and South America than GBIF, this difference in species diversity between these two databases becomes even more pronounced for the Americas (Fig 4b).

**Fig 4:**
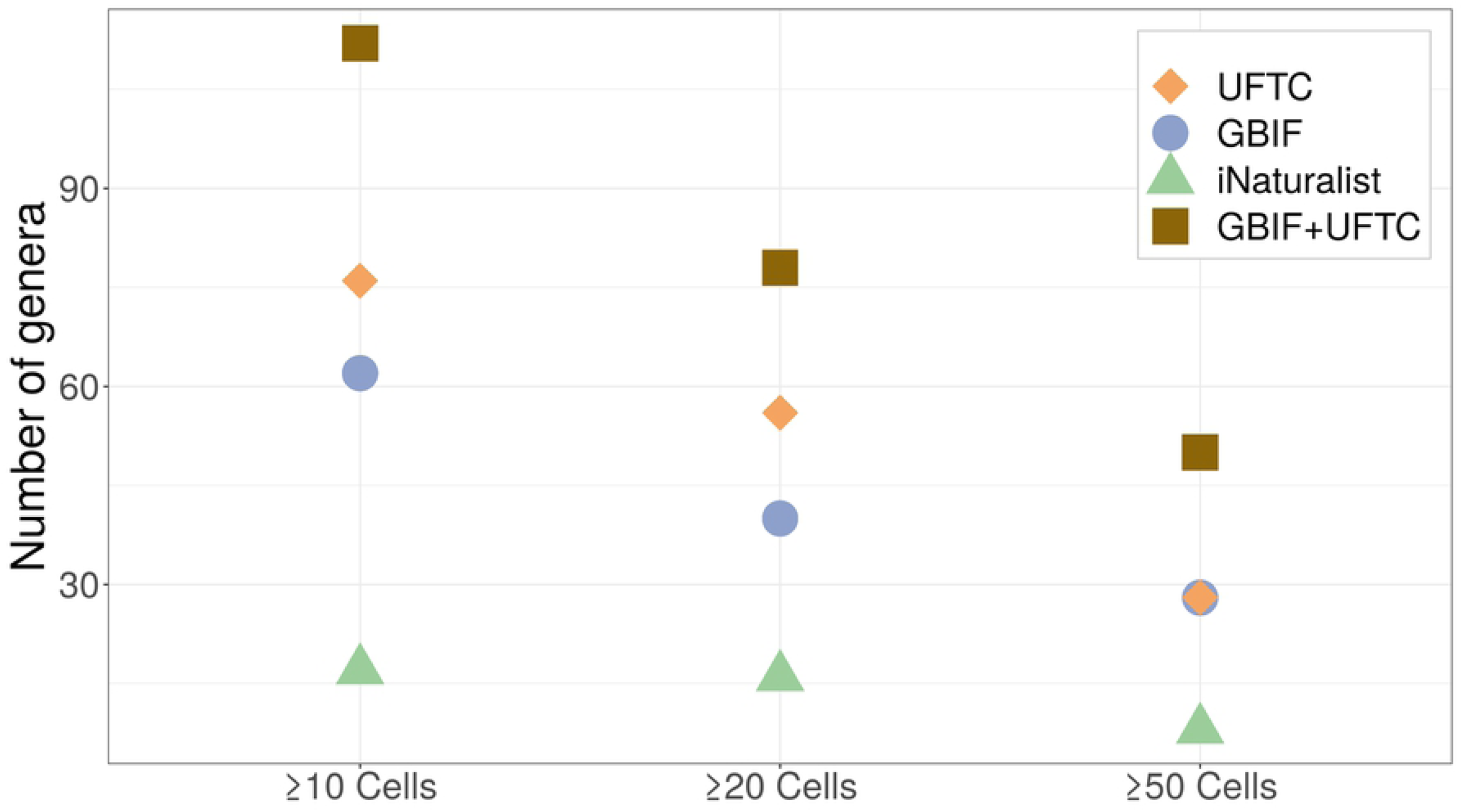
Cumulative number of termite species mapped in the UFTC, in GBIF based on expert (non-CS) observations and CS observations across the world (a) and in the Americas (b).

### Data suitability for niche modeling

The number of genera satisfying data requirements for niche modelling (may it be 10, 20, or 50 distinct cells) was largest for the UFTC, somewhat closely followed by GBIF, which in this analysis included both expert and CS data (Fig 5). Research-grade CS data played only a minor role in GBIF termite data abundance (Fig 1). As opposed to UFTC and GBIF, the iNaturalist platform (including both research grade and non-research grade) satisfied niche modeling data requirements for only few genera. Adding non-research grade observations from iNaturalist to GBIF (which itself already includes research-grade iNaturalist data) only moderately increased the number of genera meeting modeling criteria, namely by up to 14.3% for ≥ 50 distinct grid cells. As opposed to this, GBIF and UFTC appeared very complementary. A combination of these two most comprehensive sources, mostly consisting of expert and/or museum collections, led to a substantial increase in the number of termite genera suitable for niche modelling compared to either of those two sources alone (Fig 5). More specifically, complementing GBIF with UFTC data led to a stronger relative increase in genera numbers (between 78.6% and 95.0%) than complementing UFTC with GBIF data (increase between 39.3% and 78.6%). This illustrates the current dominant role of UFTC as a global termite inventory. Adding iNaturalist data as a third source to the UFTC +GBIF combination increases the number of termite genera suitable for niche modelling only slightly, namely by 1.8% (for ≥ 10 cells), 0.0% (for ≥20 cells), and 8.0% (for ≥50 cells), respectively.

**Fig 5:**
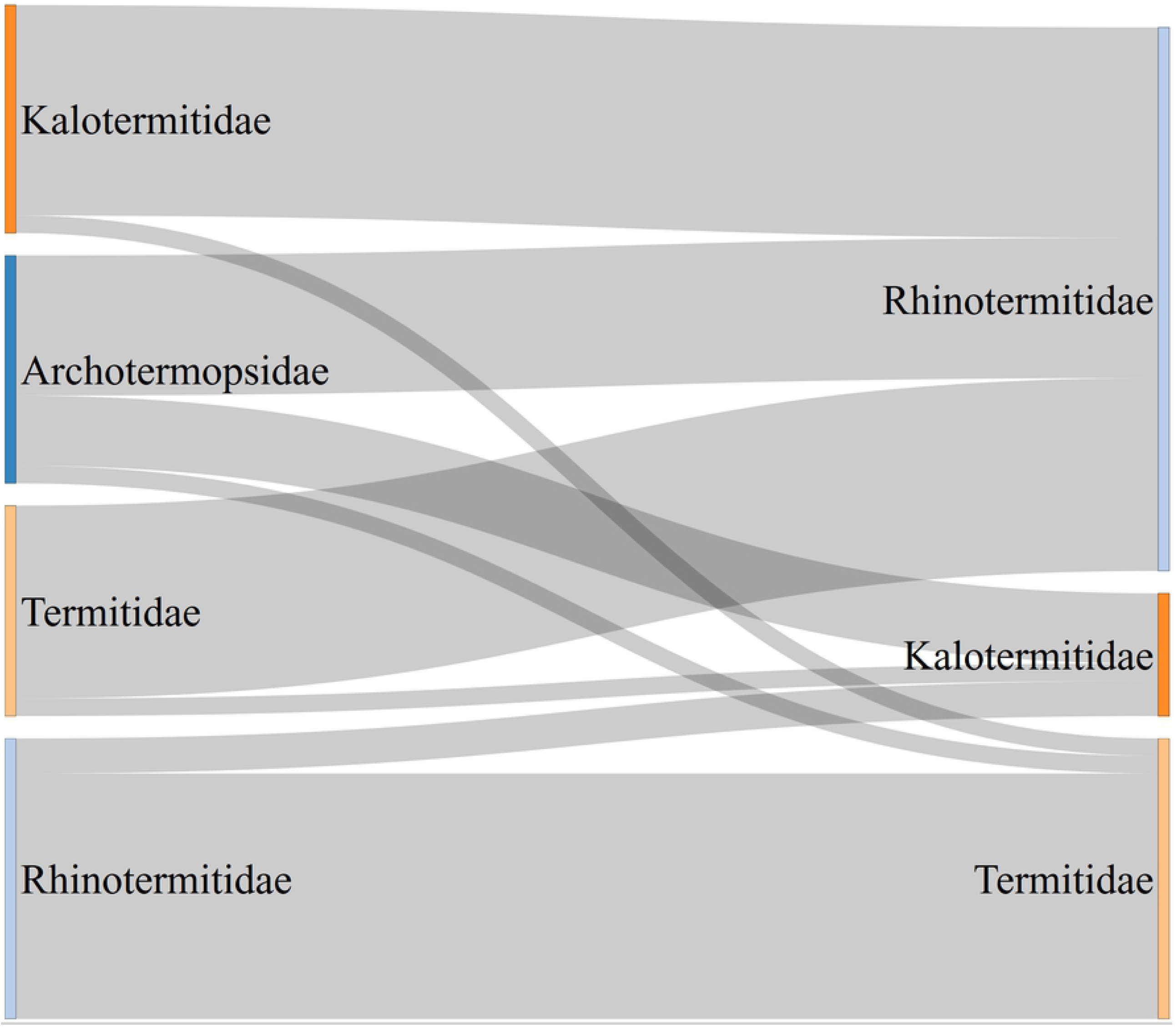
Number of termite genera covering ≥10, ≥20, and ≥50 distinct 0.1 degree cells for individual data sources and one combination.

### Taxon correctness of iNaturalist termite records

For the expert review of termite classification in iNaturalist, a subset of 2201 records that had a taxonomic description at the family level or lower was extracted (200 observations were identified at the family level, 151 at the subfamily level, 781 at the genus level, and 1069 at the species level).

The five genera of termites reported in iNaturalist which accounted for more than 72% of the observations (1597/2201) are those that are out in the open (not hidden in structures) and can thus be visually detected. These include species of genera that are very common (*Reticulitermes*, n = 939), very large (*Zootermopsis*, n = 351), build conspicuous nests (*Nasutitermes*, n = 187), or forage on the surface of the soil (*Gnathamitermes*, n = 77, and *Tenuirostritermes*, n = 43) (Table 2). All other genera had fewer than 40 observations, and many had 1 to 5 only. 351 records lack genus information but contain family or subfamily rank information only. Many photographs of the analyzed sample include the winged images which emerge from their cryptic nests during dispersal flights.

**Table 2.** Termite genera of analyzed iNaturalist records for the Americas.

| Genus | Count | Genus | Count |
| --- | --- | --- | --- |
| *Amitermes* | 10 | *Neotermes* | 3 |
| *Anoplotermes* | 12 | *Paraneotermes* | 1 |
| *Bulbitermes* | 5 | *Porotermes* | 3 |
| *Coptotermes* | 37 | *Procornitermes* | 1 |
| *Cornitermes* | 1 | *Pterotermes* | 1 |
| *Cortaritermes* | 1 | *Reticulitermes* | 939 |
| *Cryptotermes* | 3 | *Rhinotermes* | 1 |
| *Gnathamitermes* | 77 | *Rhynchotermes* | 4 |
| *Heterotermes* | 6 | *Silvestritermes* | 1 |
| *Incisitermes* | 131 | *Syntermes* | 24 |
| *Kalotermes* | 3 | *Tenuirostritermes* | 43 |
| *Microcerotermes* | 2 | *Termes* | 1 |
| *Nasutitermes* | 187 | *Velocitermes* | 1 |
| *Neocapritermes* | 1 | *Zootermopsis* | 351 |

The 2201 iNaturalist records fall into five families, which are Archotermopsidae (*n* = 351), Kalotermitidae (*n* = 198), Rhinotermitidae (*n* = 1045), Stolotermitidae (*n* = 4), and Termitidae (*n* = 603).

Most records were still in need of identification (Needs_ID quality, 79.9 %), while only 18.1 % were research grade, the remaining 2.0 % being casual observations.

68 records which did not allow for accurate identification (38 records with poor image quality, and 30 records without a picture), were removed.

Taxonomic accuracy was generally high (Table 3). That is, out of 1026 observations that were identified at the species rank in iNaturalist, 39 had an incorrect genus, 25 an incorrect family, and six an incorrect order, accounting for a total of 6.8 % observations that needed correction. Out of a sample of 1792 records for the Americas that were named at the species or genus taxonomic rank, only 78 records (4.4 %) had an incorrect genus. Conversely, irrespective of the level at which observations were identified, only 54 records out of 2133 (2.5 %) showed an incorrect family designation, and only eight records (0.4 %) showed an incorrect order, where ants or flies were mistaken as termites.

**Table 3:**
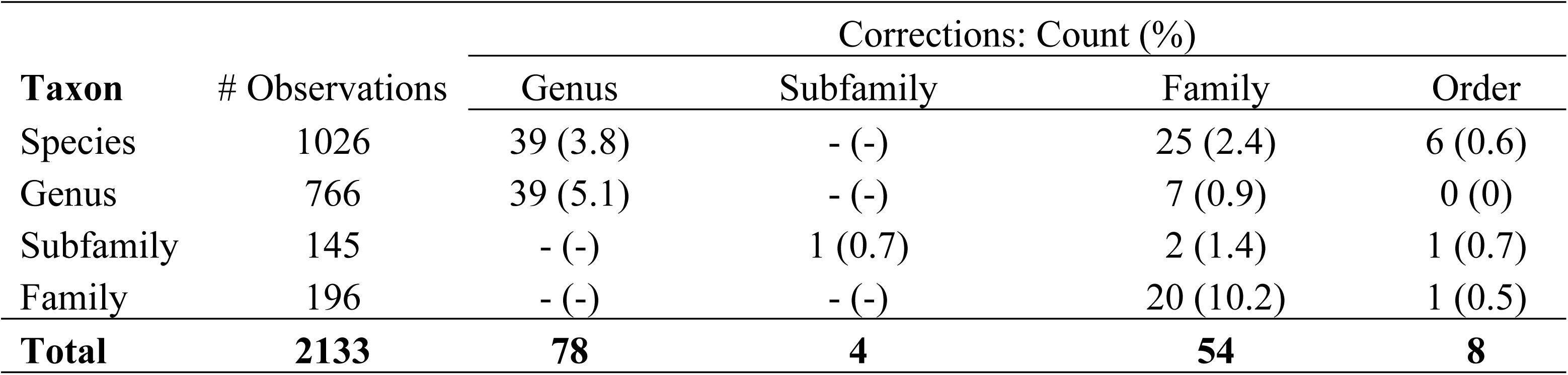
Number of observations at different taxonomic levels and the number of corrections.

| Taxon | # Observations | Corrections: Count (%) |  |  |  |
| --- | --- | --- | --- | --- | --- |
|  |  | Genus | Subfamily | Family | Order |
| Species | 1026 | 39 (3.8) | - (-) | 25 (2.4) | 6 (0.6) |
| Genus | 766 | 39 (5.1) | - (-) | 7 (0.9) | 0 (0) |
| Subfamily | 145 | - (-) | 1 (0.7) | 2 (1.4) | 1 (0.7) |
| Family | 196 | - (-) | - (-) | 20 (10.2) | 1 (0.5) |
| **Total** | **2133** | **78** | **4** | **54** | **8** |

The misclassification rates were similar between termite families (Fig 6). 13 out of 192 records (3.6 %) classified as Kalotermitidae were actually coming from other families. Corresponding numbers were 16 out of 1010 (1.6 %) for Rhinotermitidae, 12 out of 584 (2.1 %) for Termitidae, and 13 out of 343 (3.8 %) for Archotermopsidae records.

**Fig 6:**
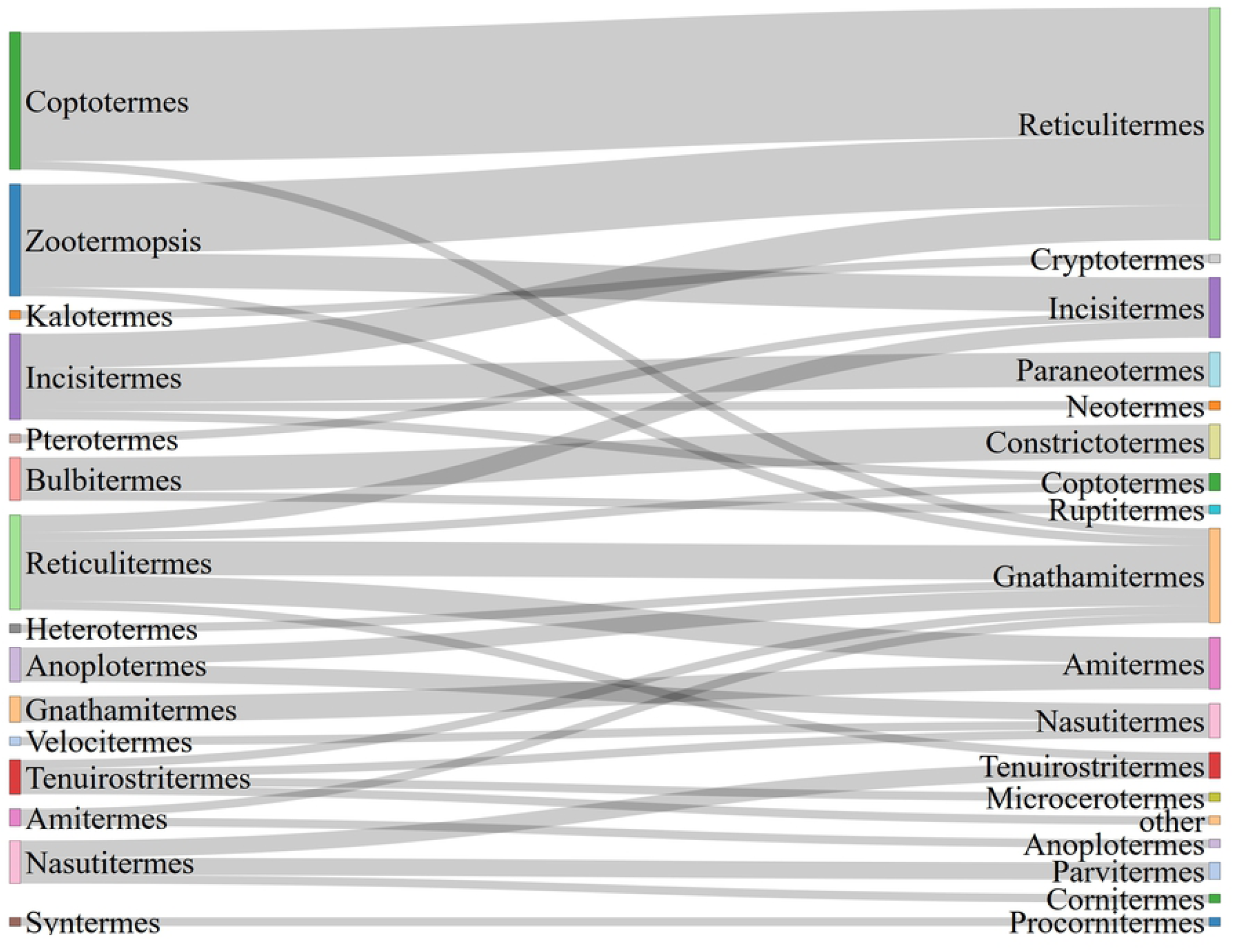
Corrections (right side) of incorrect family classifications (left side) for 54 termite records. Band-width is proportional to number of corrected observations. All four Stolotermitidae records had correct family names, which is why this family is not shown on the left side.

The same analysis at the genus level showed a more contrasted situation (Fig 7). Sixteen out of the 35 records (45.7 %) identified as genus *Coptotermes* had an incorrect genus, followed by 13/343 (3.8 %) for *Zootermopsis*, 11/906 (1.2 %) for *Reticulitermes*, and 10/906 (1.1 %) for *Incisitermes*. The high misclassification rate of the prior can likely be attributed to the fact that exotic species like *Coptotermes* are new to observers who are more familiar with native taxa like *Reticulitermes*. Superficially, both look similar.

**Fig 7:**
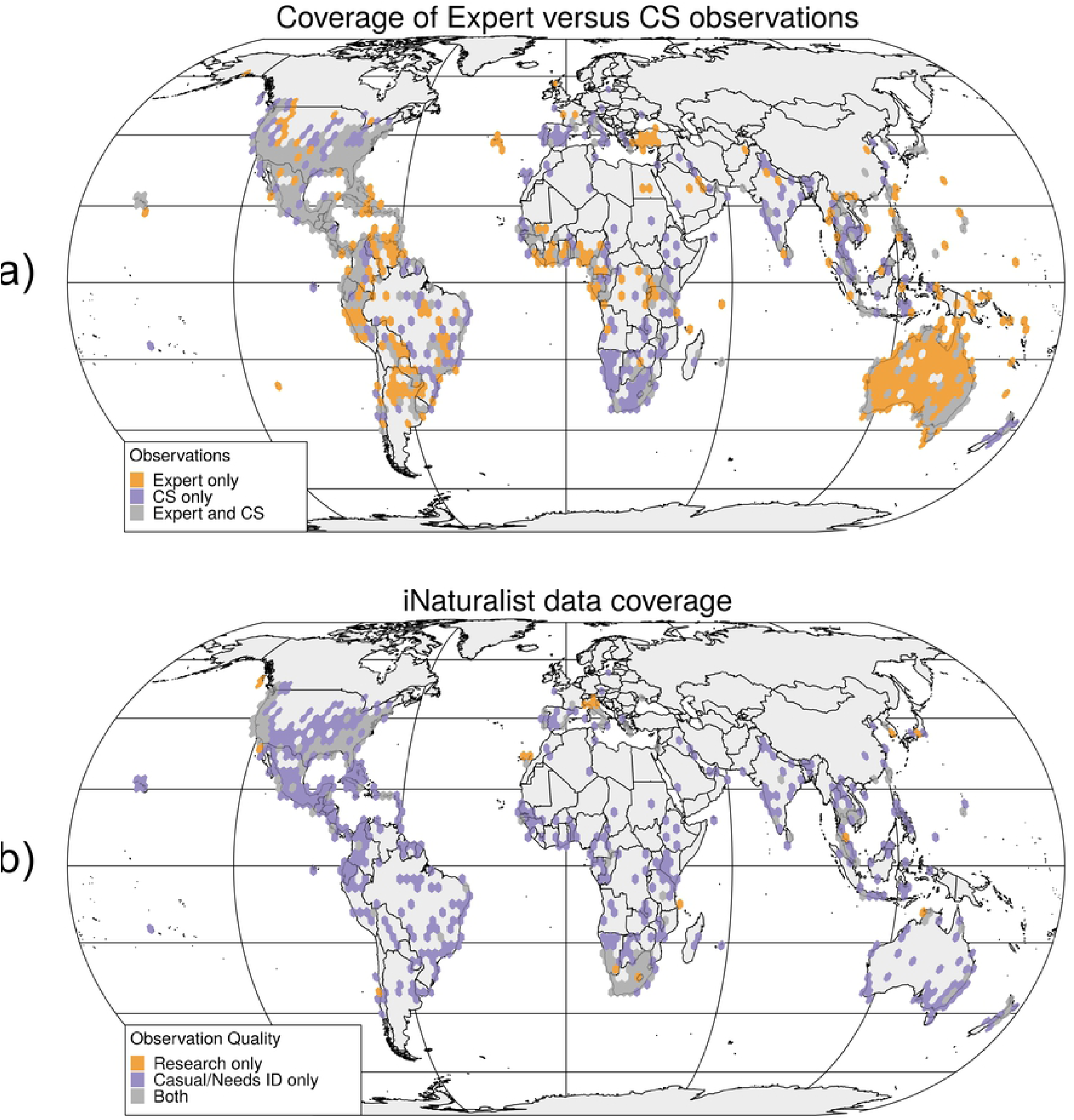
Corrections (right side) of 78 iNaturalist termite records with incorrect genus designations (left side).

Based on observation and correction counts from Table 4, a permutation independence test with 2000 randomizations showed that there was no significant association between DQA and the likelihood for a genus correction, *X*^*2*^ = 1.92, p = 0.21, and no significant association between DQA and the likelihood for a family correction, *X*^*2*^ = 2.94, p = 0.13. An explanation is that non-research grade (especially Needs-ID level data) only indicates that the necessary number of experts have not yet attempted to identify the species or genus level of an observation, but does not say anything about the accuracy of the record itself.

**Table 4.** Number of genus and family corrections for research grade and non-research grade observations based on a sample of 1792 species and genus records in the Americas.

|  | Observations |  | Genus corrections |  | Family corrections |  |
| --- | --- | --- | --- | --- | --- | --- |
|  | Count | % | Count | % | Count | % |
| Research | 394 | 22.0 | 12 | 15.4 | 3 | 9.4 |
| Non-research | 1398 | 78.0 | 66 | 84.6 | 29 | 90.6 |
| Total | 1792 | 100.0 | 78 | 100.0 | 32 | 100.0 |

## Discussion

The first part of the study showed that CS based termite observation data are still a niche product compared to expert databases and professional collections, such as museum records. Similarly, CS data did not capture much of termite species diversity. This could be because of the cryptic nature of termites and their small size, which renders field collection and species detection difficult for non-experts. Results showed that UFTC clearly outnumbered GBIF and iNaturalist in cumulative species numbers, although in the most recent years iNaturalist (and in consequence also GBIF) submissions have increased drastically, pointing towards a more important role of CS for termite observation data collection and mapping. While GBIF being a data warehouse that draws its data from numerous other data collections, it was shown that CS based collections play only a marginal role for termites (∼1.2 %) in that platform.

As of now, CS projects, global data portals that combine many data sources, as well as expert data collections, inherently come with a spatial and temporal bias, due to the opportunistic nature of CS data, the limited spatial and temporal scope of projects, and socio-economic factors limiting access to the latest technology in various parts of the world. One study, for example, compared the completeness and coverage among three open-access databases along ten taxonomic groups, finding that the coverage of systematic surveys (American Breeding Bird Survey, BBS, and federally administered fish surveys, FFS) was less biased across spatial and environmental dimensions but more biased in temporal coverage compared to GBIF data [27].

Lack of frequent updates on termite sightings can be considered a lesser problem for modeling termite species distribution, since the natural dispersal of termite colonies proceeds slowly. Alates (winged reproductives) are unable to fly more than a few hundred meters from the parent colony [28, 29]. However, occasional anthropogenic means of transportation on water, e.g. by maritime vessels [30, 31] or on land via infested ornamental timber of wood structures, allows occasional faster dispersion and establishment of new colonies.

Comparison of the world-wide contribution maps (Fig 2) showed distinct geographic biases between the three analyzed data sources. The focus of the UFTC is on the Americas since this is the primary study area of the data base creator and manager (RHS). As opposed to this, GBIF shows a strong contribution bias towards Australia through a large data collection shared by a governmental research institution (SCIRO). iNaturalist, as the only exclusively CS based platform has its geographic focus on the U.S., where the program has been founded in 2008. This shows that various factors, such as the location of individuals or that of individual organizations, as well as the country of where a project has been founded, will add to geographic contribution bias.

Analyzing the presence of genus point data across geographically distinct cells showed that the UFTC provides the highest number of termite genera that satisfy general data requirements for niche modelling, following by GBIF and iNaturalist. The combination of UFTC and GBIF increases the number of genera suitable for niche modeling by up to 95 % compared to single source data, whereas adding iNaturalist data has only a minor effect.

The third part of the study evaluated for a subset of termite records in the Americas the accuracy of record classifications in iNaturalist. Overall, it can be concluded that the classification quality of iNaturalist termite data is excellent with error rates up to only 5.1 % for genus designations and 10.2% for family designations. We found no statistically significant effect of DQA on classification accuracy.

Whether research grade records can be considered more trustworthy than non-research grade observations is important when it comes to species distribution models. This is because non-research grade termite observations in iNaturalist cover some geographic regions that are missed by research grade records (Fig 3a). Dropping non-research grade observation would therefore add to the bias of geographic coverage.

Temporal rates of submissions varied between data sources. In particular GBIF records for termites varied sporadically through the years based on punctuated submissions from professional expeditions in Mexico and Ghana. This contrasts with larger represented taxa in GBIF like Aves where professional data sets are rarely submitted and the vast majority of submissions occurs from the CS platforms [32]. While the GBIF data quality may be higher the sporadic pattern of submissions may add more noise to analysis than the steadily increasing submission rate that is frequently seen in CS. Such temporal biases can affect biodiversity models and cloud patterns when interpreting analysis [33].

## Conclusions

Using termites as an example that can be challenging to collect, we show that expert datasets, such as the UFTC and GBIF are valuable for biodiversity research because of their number of records, spatial distributions, and diversity. Especially GBIF, which taps into data from a variety of data sources, can mitigate knowledge gaps and restraints from individual projects. For termites, the contribution from CS data is much lower than expert and museum collections due to sampling biases and identification challenges. However, these CS contributions (particularly using iNaturalist’s non-research grade data set) with some extra quality controls can still be useful in biodiversity studies since they cover otherwise underexplored areas, such as central Africa or South-East Asia. Combining GBIF, UFTC, and iNaturalist data sets globally creates a more comprehensive picture of termite biodiversity than either of these data sources on their own.

Although crowd-sourced data currently play a small role in termite record numbers relative to expert or global research databases, they should not be ignored when searching for data to build termite diversity or distribution maps. More specifically, CS based data collections have the advantage of being quickly updated, and the many eyes of volunteer contributors may even be able to discover species in locations where they were not previously observed (Skejo et al. 2016). For example, in the current study, *Kalotermes approximatus* was identified as a new Texas state record from an iNaturalist photograph that was identified as an Incisitermes sp.

